# VX: an AI-enabled desktop genome viewer and transcriptome browser with a programmable analysis framework

**DOI:** 10.64898/2026.05.17.725790

**Authors:** Nikolay Shirokikh, Alice Cleynen

## Abstract

**Backgsround:** Genome and transcriptome browsers are central to the interpretation of high-throughput sequencing data, but today’s tools assume a human operator at a graphical interface and offer only limited programmability. As large-language-model assistants become routine in bioinformatics [Anthropic, 2024], this creates a bottleneck: agents cannot observe the visual state of the browser or drive it through the same interface as the human user, and analyses remain fragmented across a separate ecosystem of external tools. Transcript-coordinate data, produced by ribosome profiling [Ingolia et al., 2012] and direct RNA sequencing [Garalde et al., 2018], is also awkwardly supported in chromosome-oriented viewers.

**Results:** We present VX, a desktop genome and transcriptome viewer written in D, using GTK 3 and OpenGL, that handles genome-scale and transcriptome-scale data in a unified interface. VX exposes its full functionality through an embedded HTTP API on the loopback interface and a Model Context Protocol server of currently thirty-nine tools, so that scripts and LLM agents can load data, navigate, manage tracks, run analyses, and capture figures through the same contract used by the GUI. An integrated analysis framework provides more than fifty analyses and includes signal processing and peak calling, quantification, variant analysis, alignment statistics, interaction and cross-track comparisons, all with an explicit four-level scope hierarchy running from viewport to whole dataset; results are written to disk and, where appropriate, added as new tracks. Additional features include a magnifier popup for base-resolution inspection (Alt+hover), chromosome-alias resolution across UCSC, Ensembl, and NCBI conventions, viewport video recording via an `ffmpeg` pipe, and INI-based configuration.

**Conclusions:** VX complements existing desktop and web browsers by providing a native agent-control layer, an integrated analysis framework, and first-class transcript-space handling. The binary is freely available for non-commercial use; the HTTP API and MCP protocol are fully specified in this article, so third-party clients can be written independently of the core implementation.

## 1 Background

The interpretation of high-throughput sequencing data depends heavily on interactive visualisation. “Genome browsers”, defined as tools that render aligned reads, annotations, signal tracks, and variants against a reference coordinate system, have become essential laboratory infrastructure, and a small number of them dominate day-to-day use across the community. Web-hosted portals such as the UCSC Genome Browser [Kent et al., 2002] and Ensembl [Martin et al., 2023] provide community-curated tracks and permalink-driven navigation for hundreds of reference genomes. Desktop tools such as the Integrative Genomics Viewer (IGV) [Robinson et al., 2011, 2023] render user-supplied BAM, VCF, and signal files locally with low latency and rich per-read detail. More recently, JBrowse 2 [Diesh et al., 2023] has delivered a modular web application that scales from laptop to server and supports pluggable track types, while programmatic tools such as pyGenomeTracks [Lopez-Delisle et al., 2021] and Gviz [Hahne and Ivanek, 2016] cater to publication-quality static figures produced in scripting environments.

Two design paradigms currently dominate this landscape. The **desktop GUI** paradigm (IGV and its derivatives) prioritises responsiveness on local data; the **web platform** paradigm (UCSC, Ensembl, JBrowse 2) prioritises accessibility, shared state, and server-hosted compute. Each paradigm carries well-understood trade-offs: desktop clients are fast and private but difficult to automate; web platforms are cross-machine and shareable but add network latency and deployment complexity. A third, smaller paradigm, **script-driven static rendering**, offers reproducibility and publication quality at the cost of interactivity.

Across all three paradigms, **transcript-space visualisation** is under-served. The overwhelming majority of tools are chromosome-oriented: coordinates run along a genomic contig, and transcript structure is rendered as splice-aware overlays (for example, Sashimi plots [Katz et al., 2010]) or by collapsed exon views. Yet a growing family of experiments produces data aligned directly to transcript sequences, including ribosome profiling [Ingolia et al., 2012], long-read isoform sequencing [Amarasinghe et al., 2020], nanopore direct RNA sequencing [Garalde et al., 2018], and transcript-level quantification workflows such as Salmon [Patro et al., 2017], for which a transcript is the natural coordinate frame. Adapting genome-centric browsers to these data typically requires either rebuilding custom reference indices or rendering transcripts as synthetic chromosomes, both of which complicate provenance and confuse downstream tooling [Lauria et al., 2018, Birkeland et al., 2018].

A more recent constraint is now reshaping the analysis workflow itself. Large-language-model assistants, exemplified by tools that implement the Model Context Protocol (MCP) [Anthropic, 2024] and analogous agentic frameworks, are increasingly embedded in bioinformatics practice. Such assistants can only contribute to a visual analysis if they are able to: (a) *observe* the visual state of the viewer; (b) *drive* navigation, track management, and data loading; (c) *trigger* computations; and (d) *summarise* the results in the context of the user’s current question. Existing browsers were not designed for any of these four capabilities. Web tools expose URL schemes and, in the case of JBrowse 2, a JavaScript API, but these stop short of a full tool-use surface accessible to an external agent; desktop tools are typically controlled only through their own UI and expose no programmatic interface at all.

A secondary, long-standing pain point compounds the first: **analysis fragmentation**. Browsers visualise, while quantification, peak calling, interval arithmetic, variant annotation, and cross-track comparison are performed in separate tools (for example, BEDTools [Quinlan and Hall, 2010], deepTools [Ramírez et al., 2016], MACS [Zhang et al., 2008]). Every context switch (export, re-import, parameter re-entry) slows interpretation and discourages exploratory analyses at the point of visual inspection.

We designed VX to address these three gaps in a single application. VX is a desktop viewer that treats genome and transcriptome coordinate systems symmetrically, exposes its full functionality through an embedded HTTP API and an MCP server so that LLM agents and scripts can observe, drive, and extend sessions, and ships with a programmable analysis framework that runs ∼ 50 analyses against scoped regions of the currently loaded data. The remainder of this article describes the implementation (§2), evaluates VX against representative existing tools and a set of real-world case studies (§3), and discusses the design rationale, limitations, and future extensions (§4).

## 2 Implementation

### 2.1 Overview and architecture

VX is implemented in the D programming language [D Language Foundation], which combines systems-level performance with modern language features (Unicode handling, garbage collection, compile-time reflection, and straightforward interoperability with C). The user interface is built on GTK 3 via the gtk-d bindings [Dziemidowicz et al.], and rendering is performed using OpenGL 4.5 through the bindbc-opengl loader [Parker]. Type rasterisation uses FreeType [The FreeType Project] through a small C shim that is compiled and linked during the build. The project is assembled with the standard D build tool *dub*, which provides separate configurations for Linux (primary), macOS, and Windows cross-compilation. The Linux configuration ships four release profiles tuned to different CPU baselines (`release-aggressive` for x86-64-v3 broad-compatibility, `release-znver2` and `release-broadwell` for the two most common workstation microarchitectures, and `release-native` for build-host-exact tuning); all four share full LTO with static druntime+phobos and produce a self-contained Linux binary of approximately 19 MB. The `release-windows` (MSVC-linked, x86-64-v3, bundled GTK 3 DLLs) and the macOS `.app` bundle (arm64, ad-hoc signed, `dylibbundler`-bundled dependencies) are produced from the same source tree without source-level changes.

The source tree is organised into eight top-level modules that correspond to wellseparated concerns: `data` (parsers and data structures for FASTA, GTF/GFF, BAM, bigWig, bigBed, VCF), `tracks` (domain types for each visualisable track category), `visualiz--ation` (the OpenGL renderer, font atlas, and magnifier), `ui` (the GTK main window, control panel, settings, and dialogues), `net` (the embedded HTTP API server), `analysis` (the 50+ built-in analyses and their execution engine), `config` (INI-based persistent configuration), and `recording` (the `ffmpeg`-backed video recorder). An illustration of the different modules is shown if Figure 1 while a schematic of their dependency relations is shown in Figure 2.

**Figure 1:**
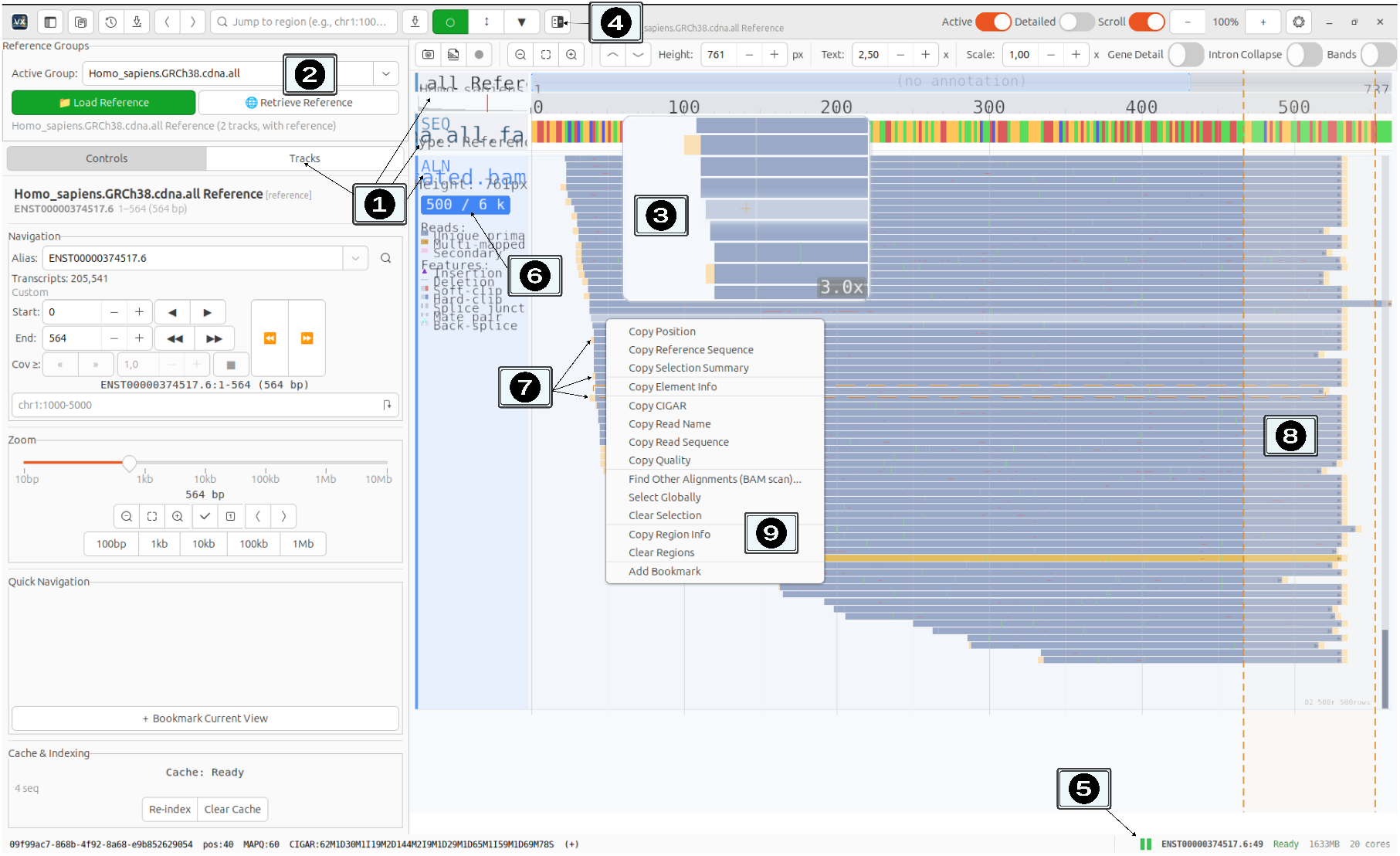
VX human interface. Example on a human transcriptome-aligned dataset. **1:** reference, sequence and alignment (bam) tracks are displayed. Their display and visualisation of additional tracks can be controlled via the “tracks” button. **2:** groups can be selected in the active group tab, allowing the user to switch from groups of references and tracks to other groups without reloading anything. **3:** the magnifier popup (Alt+hover) provides a base-resolution view anchored to the cursor without leaving the viewport. **4:** The Analyse button exposes the analyses applicable to the currently selected or visible tracks. **5:** the load-status strip indicates the loading status of each track, warning users when data has not yet been pulled from disk. **6:** the subsampling badge indicates when read-subsampling is performed to visualize the track. Level of subsampling can be selected independently for each track in the track control tab. **7:** soft-clip visualization as a yellow strip matching soft-clip length. **8:** zooming in window can be selected with the shift controls to investigate regions of interest. **9:** right-click on a read allows user to retrieve read-specific features, including sequence, CIGAR, or alternative alignment of the read.

**Figure 2:**
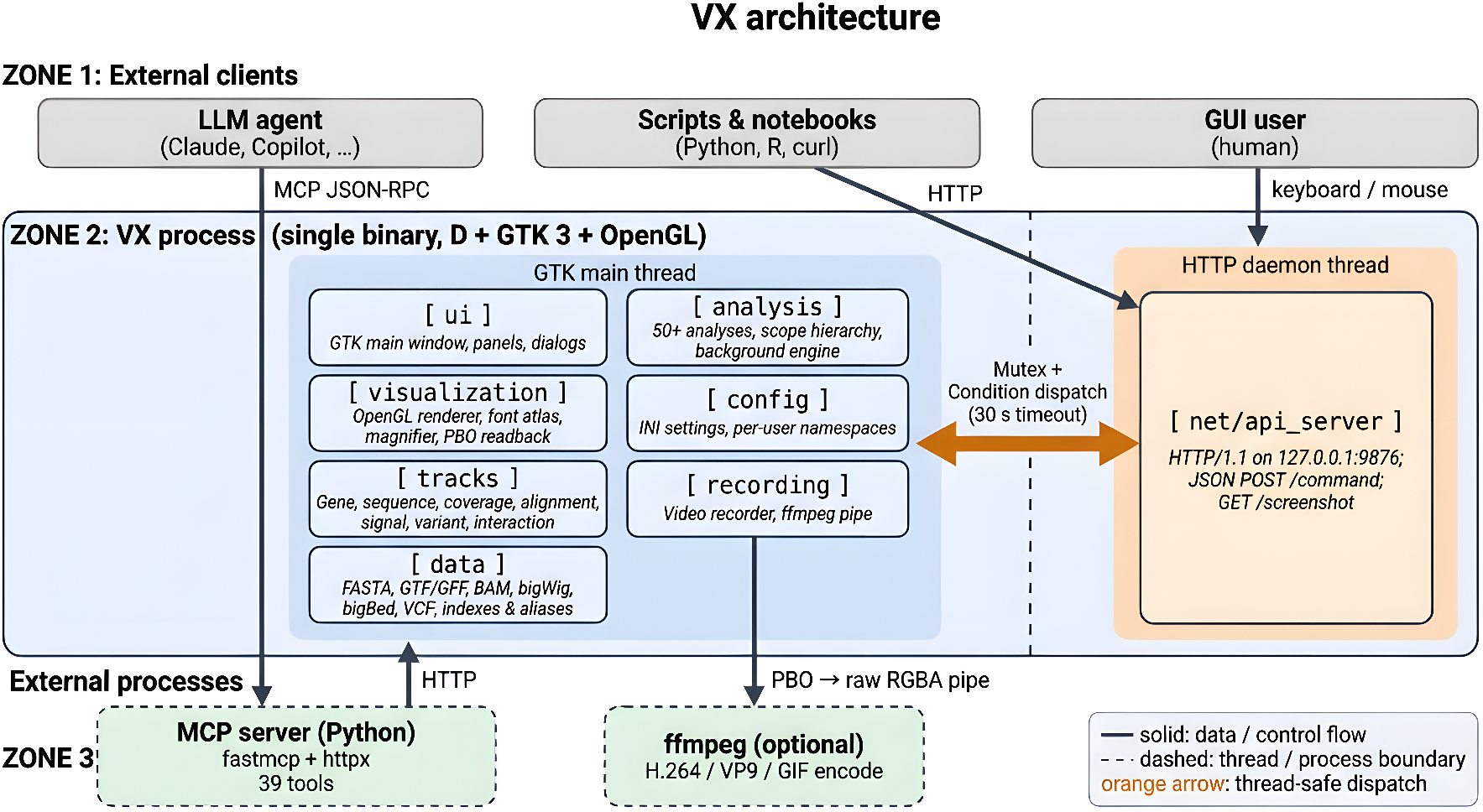
VX architecture. The D core (`data, tracks, visualization, ui, analysis, config, recording`) is driven by the GTK main thread. The embedded HTTP server (`net`) runs on a separate daemon thread and marshals external requests onto the main thread via a `Mutex/Condition` pair. The Python MCP server is a thin wrapper over the HTTP API and is launched as a separate process by MCP-aware clients. Optional external dependencies (`ffmpeg`) are engaged only when needed.

The VX process is single-binary: no external service is required for the application itself. Only two subsystems interact with external processes: the optional `ffmpeg` pipe used for video recording, and the optional Python MCP server described in §2.5, which is launched as a separate process by MCP-aware clients. Configuration is persisted to an INI file next to the executable; nothing is written to system locations, and a fresh working directory produces a reproducible default state.

### 2.2 Data model and track system

VX organises visualisable data into *tracks* and *groups*. A track is a typed collection of features or signal values defined over one or more contigs; the track system currently supports gene (GTF/GFF-derived), sequence (FASTA), coverage (BAM-derived), alignment (BAM reads at read level), signal (bigWig), variant (VCF), interaction (contact-matrix formats), and generic BED/bigBed tracks. Each track type implements a common rendering and interaction interface but retains its own file reader, feature model, and drawing routine, so new track types can be added without touching the shared code paths. The rendered appearance of the supported track categories is illustrated in Figure 1, block 1.

A group bundles a reference sequence together with the tracks that share its coordinate system (Figure 1, block 2). This keeps multi-reference sessions organised: a user may have one group holding a human GRCh38 genome with its annotations and alignments, and a second group holding a *Saccharomyces cerevisiae* assembly with its own annotations, and switch between them without contaminating either. The group abstraction also underlies the transcriptome browsing mode: because VX treats any named contig identically, a FASTA containing transcripts (for example, Ensembl ENST identifiers) with a matching transcript-level GTF and a transcriptome-aligned BAM produces a fully functional transcript browser without any code changes (§3.6). The same rendering, analysis, and navigation machinery applies.

Cross-convention naming is handled by a dedicated alias resolver that translates between the three major chromosome naming conventions (UCSC `chr1`, Ensembl `1`, NCBI RefSeq `NC_000001.11`) at the API and query layers. This means a user can load a reference in one convention and annotations in another, and VX will reconcile them transparently.

To keep startup and navigation responsive on genome-scale data, the parsers are all streaming or region-indexed: the GTF parser builds an interval index on first load, BAM reads are fetched by genomic region via the standard BAM index [Li et al., 2009], which can be created on the fly, and bigWig and bigBed data are queried through their native random-access structures [Kent et al., 2010]. Large files are never loaded into memory in full.

### 2.3 Rendering pipeline

The rendering pipeline is what gives VX the user-facing properties on which the rest of the application depends: smooth pan and zoom across many orders of magnitude (from a single base pair up to a whole chromosome) while displaying multiple tracks simultaneously, HiDPI-correct text and gene glyphs, an interactive in-viewport *magnifier* popup for inspecting sub-pixel detail without zooming the whole view, and direct video capture of the rendered scene for figure and supplementary-movie production. The remainder of this subsection describes the implementation choices that make those capabilities tractable; a reader interested only in the user-facing behaviour can safely skip to §2.4.

All viewport drawing in VX is performed through a single OpenGL context attached to a GTK `GLArea` widget. The renderer targets OpenGL 3.3 core and uses a small set of retained vertex-buffer objects rather than legacy immediate-mode calls, so that scrolling and zooming at interactive rates remain feasible even with many tracks on screen. An orthographic projection matches one OpenGL unit to one logical pixel; physical-pixel conversion for HiDPI displays is applied separately inside scissor and viewport calls, so track layout code can remain resolution-independent.

Text is rasterised by a FreeType-backed font atlas (`visualization/font_atlas.d`). FreeType is accessed through a short C shim that wraps the parts of its API used by VX, which avoids binding-version fragility and keeps the D side free of compile-time C types. The atlas supports configurable font families, an extended Unicode range, and hot re-initialisation when the user changes font settings at runtime.

To keep rendering bounded when a user zooms out, tracks delegate to a **level-of-detail** (LOD) renderer that selects among four regimes (*Density, Boundaries, Features*, and *Sequence*) according to the current base-pairs-per-pixel ratio. At the coarsest regime a track draws a density summary rather than individual features; at the finest, the raw DNA sequence is drawn glyph-by-glyph. The selection is continuous, so a track smoothly transitions between representations as the user zooms.

The **magnifier popup** (a novel UX element) is implemented as a second render pass into the same framebuffer. When the user holds Alt and hovers, the renderer reconfigures the projection matrix to a sub-region of the scene (`updateProjectionMatrix(srcW, srcH, srcX, srcY)` in `visualization/gl_renderer.d`), enables `GL_SCISSOR_TEST` clipped to the popup rectangle, and re-issues the draw calls for the affected tracks. The result is a magnified view rendered using the same geometry as the main viewport, not an image upscale, so lines, glyphs, and gene features remain sharp at any magnification (1.5*×* –10*×*, configurable) as shown in Figure 1, block 3.

**Video recording** is implemented as a second consumer of the same rendered frames. After each `onGLRender` completes, the `VideoRecorder` in `recording/video_recorder.d` issues an asynchronous `glReadPixels` into one of two pixel-buffer objects (PBOs), and maps the other PBO (which holds the previous frame) for reading by the CPU. The CPU-side pixels are flipped top-down and written as raw RGBA to the standard input of a spawned `ffmpeg` [FFmpeg Developers] process, which encodes the stream directly into MP4 (H.264), WebM (VP9), or GIF. Double-buffering the PBOs hides the GPU-to-CPU transfer latency behind the next frame’s rendering, and piping raw frames to `ffmpeg` avoids any intermediate disk encoding step.

### 2.4 Embedded HTTP API server

The HTTP API turns VX into a scriptable instrument. From any language that can issue an HTTP request, an external process can load files, navigate the viewport, toggle track visibility, run analyses, capture screenshots, and control video recording. In essence, the same surface a human user drives through the GUI is exposed as a single stable network endpoint. This is what enables reproducible figure scripts, batch capture for supplementary movies, remote-controlled demonstrations, and (via the MCP wrapper of §2.5) LLM-driven exploratory sessions. The remainder of this subsection describes how that surface is implemented and what guarantees it provides.

The defining architectural choice in VX is that every user-visible action (loading a file, moving the viewport, changing a track’s visibility, running an analysis, capturing a screenshot) is reachable through a programmatic interface. This is implemented as a small HTTP/1.1 server that is embedded directly in the VX process and bound to the loopback interface at `127.0.0.1:9876` by default (port and bind address are configurable). The server is implemented in D using the standard library’s socket primitives and runs on a dedicated daemon thread so that the GTK main loop is never blocked by network I/O.

The protocol is deliberately minimal. Two endpoint families cover the entire feature surface:

- `POST /command` accepts a JSON body of the form `{“command”: “<name>“, “params” : { … }}` and returns a JSON response. Error responses include both a human-readable `error` field and a stable machine-readable `code` (for example INVALID_PARAMS, `NOT_FOUND`, `NOT_READY`), which clients can use for reliable dispatch.
- `GET /screenshot` and `GET /screenshot/viewport` return PNG-encoded images of the full window or the rendering viewport respectively, with an optional scale factor for higher-resolution capture.

Because every VX command must ultimately touch GTK widgets or the OpenGL context, each incoming HTTP request is marshalled from the network thread onto the GTK main thread using a pair consisting of a Mutex and a Condition variable. The network handler submits the work via `g_idle_add`, then blocks on the condition variable until the main thread completes the operation and publishes the result. A configurable dispatch timeout (30 s by default, bounded between 5 s and 120 s) protects the network thread from a stalled main thread. This pattern preserves the thread-safety contract of GTK and OpenGL while giving external clients a fully synchronous request/response interface.

The embedded-server design has several practical consequences. First, because the server is loopback-only by default, no authentication is required for the common single-user scenario, and there is no attack surface exposed to the wider network. Second, any language that can speak HTTP (*e*.*g*. Python, R, Julia, JavaScript, command-line tools such as `curl`) becomes a first-class VX client. Third, the server presents a stable interface contract that is independent of the internal D implementation; this enables the third-party tooling strategy discussed in §4.2.

### 2.5 Model Context Protocol server

Large-language-model assistants increasingly drive exploratory bioinformatics workflows [Thirunavukarasu et al., 2023]. The Model Context Protocol (MCP) [Anthropic, 2024] is an open specification developed to let such assistants communicate with external tools through a uniform JSON-RPC-style interface. VX provides an MCP server as a thin wrapper around the HTTP API described in §2.4. The server is implemented in Python using the `fastmcp` framework [Lowin] and the `httpx` client library [Christie et al.], and is distributed alongside the application.

The wrapper exposes 39 tools (currently), grouped by task category (Table 2). Each MCP tool corresponds to either a single HTTP endpoint or a small composition of endpoints, and each returns structured output that is directly usable by an LLM client. This design produces a sharp separation of concerns. The D core is responsible for correctness, performance, and visual fidelity; it exposes a stable wire-level contract. The Python MCP server translates that contract into the MCP tool vocabulary, handles connection errors, and shapes responses for LLM consumption. Because the MCP server is published in full, a third-party developer can extend it with custom tools, wrap it for a different MCP client, or replace it entirely with a bespoke implementation in another language. An analogous server could be written for any future agent protocol without touching the D core.

**Table 1:**
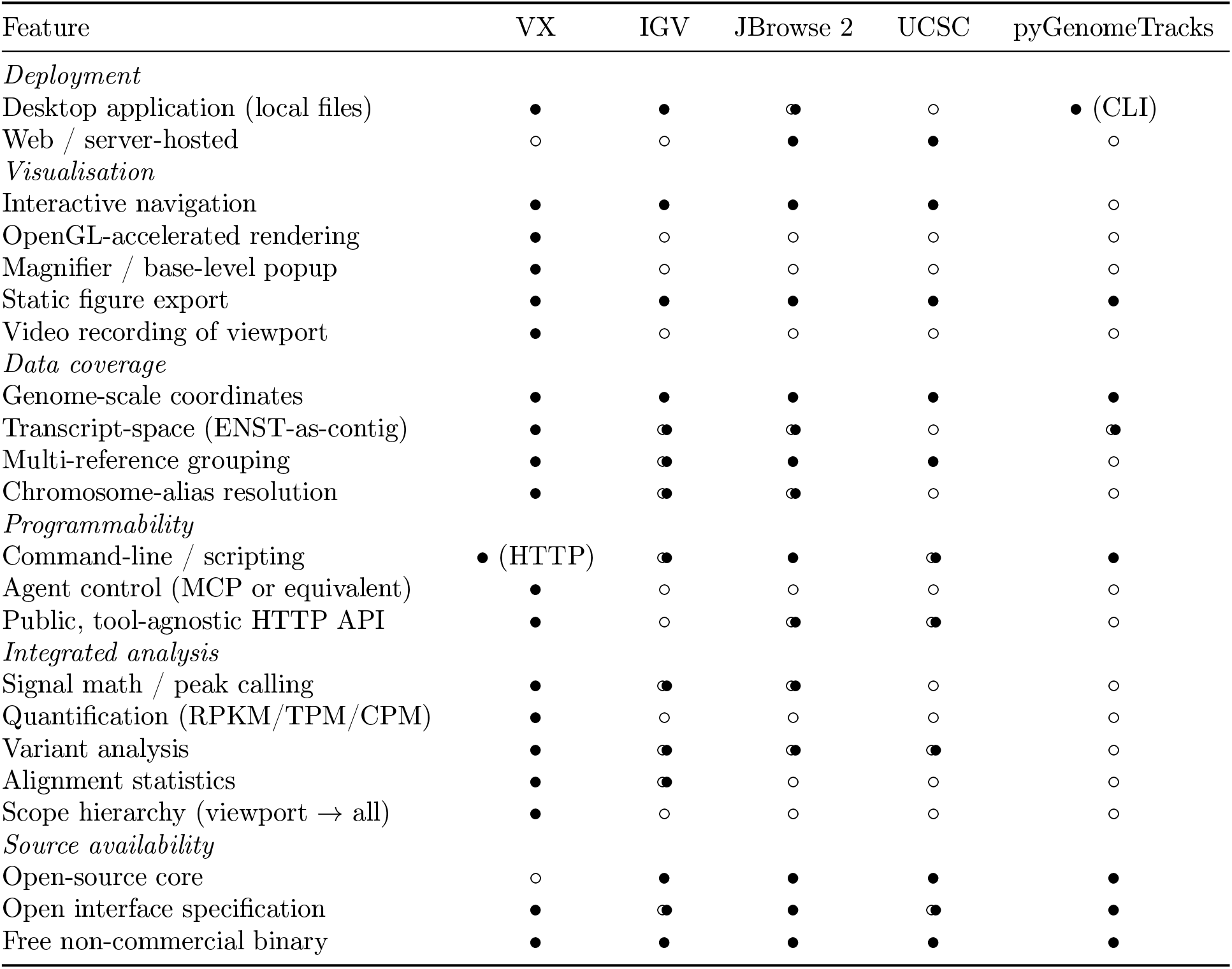
Feature comparison of VX with representative genome-visualisation tools. Marks indicate the nature of support: • native, ◦• partial or via external tooling, ◦absent; they are not a qualitative ranking.

**Table 2:** MCP tool categories exposed by VX (39 tools total).

| Category | *n* | Tools |
| --- | --- | --- |
| Observation | 5 | vx_ping, vx_get_state, vx_screenshot, vx_screenshot_track, vx_list_chromosomes |
| Navigation | 4 | vx_navigate, vx_zoom, vx_pan, vx_select_region |
| Data loading | 1 | vx_load_file |
| Track management | 4 | vx_set_track_visibility, vx_set_track_height, vx_remove_track, vx_reorder_track |
| Group management | 3 | vx_create_group, vx_switch_group, vx_remove_group |
| Track-type options | 4 | vx_set_alignment_options, vx_set_signal_options, vx_set_variant_options, vx_set_interaction_options |
| Data queries | 3 | vx_get_sequence, vx_get_genes, vx_search_genes |
| Bookmarks | 4 | vx_add_bookmark, vx_list_bookmarks, vx_remove_bookmark, vx_goto_bookmark |
| Loading status | 2 | vx_loading_status, vx_cancel_loading |
| Export | 3 | vx_export_region, vx_export_svg, vx_svg_export_status |
| Configuration | 1 | vx_set_config |
| Analysis framework | 4 | vx_list_analyses, vx_run_analysis, vx_analysis_status, vx_cancel_analysis |
| Video recording | 3 | vx_start_recording, vx_stop_recording, vx_recording_status |
| **Total** | **39** |  |

### 2.6 Integrated analysis framework

The second pillar of VX’s design is that common computational-biology analyses are built into the application rather than deferred to external pipelines. In a conventional workflow, even routine operations such as computing GC content for a region, intersecting two BED files, deriving a log-ratio signal from two bigWig tracks, or calling peaks above a background input, require leaving the viewer, invoking a command-line tool, writing the output back to disk, and re-loading it as a new track. Each of those hops adds friction, breaks the visual feedback loop, and is a barrier to agent-driven exploration because the agent must reason about an external toolchain, file-system layout, and result-tracking conventions. VX collapses that loop into a single in-process call. We treat this not merely as a convenience layer but as a first-class architectural contribution, organised along four design axes: (i) a uniform on-demand **scope hierarchy** that exposes the same analysis at viewport, chromosome, group, and whole-corpus granularity; (ii) a **result-as-track pipeline** that re-ingests outputs as renderable tracks so compositions chain naturally; (iii) breadth of **built-in catalogue** that subsumes the operations users would otherwise script ad hoc; and (iv) full **MCP exposure** so every analysis can be invoked, parameterised, and monitored by an LLM agent with no manual scripting.

#### Catalogue breadth

The framework currently comprises more than fifty analyses organised into seven functional families accessible via a single click (see Figure 1, block 4): **signal mathematics** (signal difference, ratio, *Z*-score normalisation, peak calling, enrichment over input, prominence, gap detection), **sequence analysis** (GC content, nucleotide frequency, CpG density, motif search, repeat density), **track operations** (interval intersection, subtraction, union, closest-feature mapping), **quantification** (feature counts, RPKM, TPM, CPM, per-exon coverage, bin statistics), **variant analysis** (annotation, Ts/Tv ratio, density, allele-frequency spectrum, intersection, coding impact), **alignment statistics** (mapping quality, insert size, strand bias, duplicate rate, soft-clipping, chimeric reads, read length, per-base mismatch rate), **interaction analysis** (contact frequency versus distance, enrichment, TAD-boundary detection), and **cross-track** methods (Pearson correlation, signal at features, heat-map matrix generation, profile plots). The set was chosen by surveying the operations that recur across the workflows that VX is intended to support (RNA-seq exploration, variant inspection, transcriptome browsing, long-read inspection, contact-map analysis), and by ensuring that each major track type has a nontrivial analysis family targeting it; coverage is therefore intentionally horizontal rather than depth-first within a single domain.

#### On-demand scope hierarchy

A single design decision pervades the analysis surface: every analysis accepts the same four-level `scope` parameter, regardless of input type or output format. `viewport` restricts the computation to the currently visible genomic region; `chromosome` (or, in transcriptome mode, `transcript`) runs it over the contig containing the viewport; `group` runs it over every track in the current reference group; and `all` runs it over every loaded track across every group. The same scope token applies whether the user is computing GC content, intersecting BED intervals, or calling peaks on a bigWig signal. This orthogonality is what makes the framework usable during exploration: a user can run a quick viewport-scope iteration to inspect a hypothesis, then promote the same call to chromosome or group scope when they want a publishable summary, with no re-parameterisation. In agent-driven workflows the gain is sharper still: an LLM can cheaply iterate viewport-scope analyses while forming a hypothesis and only commit to a genome-scale run once the question has narrowed (case study 1, §3.4).

#### Result-as-track pipeline

Every analysis that produces a spatially indexable output (intervals, signal, variants, contacts) writes the result to a user-configurable directory in a standard format (BED, BedGraph, VCF, or TSV, as appropriate to the analysis type) and, in the same atomic step, registers the file back into the current group as a new track of the appropriate type. The newly-created track participates in rendering, selection, export, and, crucially, in further analyses without any additional user action. Pipelines compose naturally as a result: the output of an interval-intersection becomes a valid input to feature counting; the output of a peak call becomes a valid input to GC-content computation in those peak regions; the output of a *Z*-score normalisation becomes a valid input to cross-track correlation. The same composition holds whether the chain is invoked from the GUI, the HTTP API, or an MCP client, because the registration step is internal to the analysis engine rather than living in a workflow layer above it.

#### Execution and cancellation

All analyses run on a dedicated background thread managed by a singleton engine, leaving the GTK main loop responsive to user input during computation. Progress is reported through a non-blocking callback channel that the GUI renders as a per-analysis bar in the status strip and that the HTTP/MCP layer surfaces through `vx_analysis_status`. Any analysis, including long-running chromosome-or group-scope runs, can be cancelled cleanly through the same control surface that launched it (GUI button, `vx_cancel_analysis` MCP call, or HTTP `POST /command`); cancellation is cooperative and deletes the partial output file rather than leaving a half-written track registered in the group.

#### MCP exposure and agent-driven analysis

The analysis framework is fully addressable from the Model Context Protocol layer(§2.5) through four cooperating tools: `vx_list_analyses` returns the catalogue filtered by a supplied set of input tracks (so an agent can ask “what can I run on this BAM and this bigWig?” and receive a typed list of applicable analyses with their parameter schemas); `vx_run_analysis` accepts the analysis name, the track set, a `params` object including the `scope` token, and an optional output path, and returns immediately with a job handle; `vx_analysis_status` returns the current progress, intermediate diagnostics, and (on completion) the registered output track identifier; `vx_cancel_analysis` terminates a running job. Combined with the result-as-track pipeline, this gives an LLM agent the ability to plan and execute multistep analytical pipelines without ever leaving the protocol boundary: the agent can list applicable analyses, dispatch one, wait on its status, observe the new track it produced via `vx_get_state`, take a screenshot via `vx_screenshot`, and dispatch the next analysis that takes the new track as input. To our knowledge no other genome viewer currently exposes a comparable catalogue through an agent-protocol interface; the case studies in §3 demonstrate this end-to-end loop in practice.

### 2.7 Responsive feedback during long operations

A recurring constraint in genome browsers is that some operations cannot be made imperceptibly fast: opening a 23 GB BAM, building a per-contig coverage summary, or running a chromosome-scope analysis all take time that the user will feel. VX treats this as a design problem rather than an inevitability: whenever an operation cannot complete inside a single render frame, the application surfaces what it is doing, what stage it is at, and (where known) what proportion remains, so that the user can distinguish a slow but healthy operation from a stalled one. Two concrete manifestations of this principle are the load-status strip and the alignment-track subsampling badge.

The **load-status strip** (see Figure 1 block 5) is a one-line widget anchored beneath the control panel that aggregates two classes of long operation: the initial *file-open phase* (reference indexing, GTF parse, BAM index probe) and the per-region *read-fetch phase* that fires when the user navigates into a window for which alignment data has not yet been pulled from disk. The strip names the file or region, the current stage, and an elapsed counter; on completion it briefly shows a success summary before fading. The widget is driven by the same progress channel that the analysis engine already uses (§2.6), so a long analysis, a long load, and a long region-fetch all show up in a single consistent surface; the HTTP/MCP layer exposes the same state through `vx_loading_status` (and, for analyses, `vx_analysis_status`).

The **subsampling badge** (see Figure 1 block 6) addresses a more subtle case. When an alignment track resolves a viewport whose read count exceeds the per-frame cap (`max_reads`, default 500; configurable per session and per-track, alongside a `min_mapq` filter), VX draws a subset of reads selected uniformly at random rather than blocking the render or silently dropping data. To prevent that subsampling from being misread as a coverage drop, the track header shows a small badge stating that the view is sub-sampled and what fraction of the total reads it represents. The same cap is honoured by the screenshot endpoint, so any figure produced through the HTTP API carries the same subsampling semantics (and the same badge) as the interactive viewport, eliminating a class of silent figure artefacts that we encountered repeatedly in early testing on large short-read datasets. The cooperating subsystems that enable the strip-and-badge UX are asynchronous BAM read-fetching off the GTK main thread, idle-callback marshalling of progress updates back to the GUI, and a single non-blocking progress channel shared by load, fetch, analysis, video record, and SVG export. Together, they implement the underlying rule that any operation longer than a render frame must declare itself.

Pre-release testing on a 23 GB BAM informed the design of both features: without them, a long first-region fetch was indistinguishable from a hang, and a subsampled alignment view was indistinguishable from a low-coverage one.

### 2.8 Configuration and persistence

VX persists all user-facing settings in a single INI file, `genome-viewer.cfg`, placed alongside the executable. The INI format was chosen over JSON or YAML because section-oriented plain text is directly editable by users and by scripts, survives diff and merge cleanly, and carries no schema dependency. The configuration covers window geometry and interface scaling, HiDPI and font settings, path defaults (references, results, snapshots, sessions, tracks), view and display preferences, rendering performance parameters (cache size, loader threads, LOD toggle, BAM read caps), selection and status-bar styling, image-export and viewport settings, analysis defaults, startup behaviour, MCP/API bind address and dispatch timeout, video-recording presets, per-track visual styles, font configuration, and keyboard hotkeys. Each concern has its own `[Section]` so users can edit a single area without touching the rest. The loader also supports per-user namespacing (`[user.Section]`) on shared machines and falls back to compiled-in defaults for any missing key, which guarantees that an older configuration file remains valid after an upgrade.

### 2.9 Use of large language models in development

In line with the journal’s policy on artificial-intelligence tooling, we disclose that Claude Code (Anthropic) was used as a coding assistant during the development of VX. Its role was circumscribed to generating and reviewing source-level implementation details, for example scaffolding parser routines, proposing refactors, and drafting unit-level boilerplate, under continuous human direction. All scientific claims, architectural decisions, benchmark designs, and the prose of this manuscript are the work of the human authors, who take full responsibility for the content. No patient data, unpublished third-party code, or confidential material was provided to the assistant. The VX binary and the published HTTP and MCP interface specifications were compiled, run, and audited independently of the assistant.

## 3 Results

### 3.1 Feature overview

A typical VX session illustrates how the components of §2 combine at the user level (Figure 1). On launch, VX presents an empty viewport, a left-hand control panel, and a header bar with a search entry and an *Analyse* button. The user opens a FASTA reference, at which point VX creates a group keyed to that reference and populates the contig list. A matching GTF or GFF added to the same group yields a gene track with exons, introns, and transcript structure; a BAM file is then loaded as a coverage track, an alignment track, or both.

Navigation proceeds through gene-name search, coordinate entry, click-and-drag panning, and keyboard shortcuts, supplemented by the magnifier popup (Alt+hover), which provides a base-resolution view anchored to the cursor without leaving the viewport. The *Analyse* button exposes the analyses applicable to the currently selected or visible tracks; a chosen analysis runs on a background thread with its scope (viewport, chromosome, group, or whole dataset) selected at launch, and its results are written to disk and, where appropriate, added as new tracks. The same session is reachable through the HTTP API and the MCP server (§2.4, §2.5), so an LLM agent can drive the same workflow, capture screenshots, and summarise the resulting state. Figures for the manuscript itself are produced either through the built-in static export or through the PBO-based video recorder described in §2.3.

### 3.2 Feature comparison

The feature matrix (Table 1) summarises how VX sits relative to four widely used comparators. IGV [Robinson et al., 2011] remains the reference desktop browser, with broad format support and interactive responsiveness on local files; JBrowse 2 [Diesh et al., 2023] is the leading modern web platform, with a mature plugin ecosystem; the UCSC Genome Browser [Kent et al., 2002] is the canonical reference for large-scale comparative genomics delivered over the web; pyGenomeTracks [Lopez-Delisle et al., 2021] is a widely used command-line tool for reproducible static figures. Each has shaped the field and continues to serve workflows for which it was designed.

VX does not aim to replace any of these. Its contribution is concentrated in three rows of the matrix: agent control via MCP, a tool-agnostic public HTTP API exposing the full feature surface, and a built-in analysis framework with an explicit scope hierarchy. Two further rows, native transcript-space support and video recording, extend the desktop browsing paradigm in directions that complement the existing tools rather than compete with them. The trade-off is that VX’s core is not currently open-source; we address this directly by publishing the full HTTP and MCP interface specifications, which together constitute a sufficient contract for third-party clients, scripts, and bindings to be written independently of the D implementation.

### 3.3 Performance benchmarks

We characterised VX on a commodity desktop (AMD Ryzen 5 4600G, Zen 2 6 cores / 12 threads, 30 GiB RAM, integrated Vega 7 graphics, btrfs home, Manjaro 6.19.8) against IGV 2.19.7 (Java 21, `-Xmx10.5 GB`) on the same machine with identical inputs: the full ENCFF754JEN BAM (23.2 GB, 3.15 *×* 10^8^ reads, GRCh38), the GRCh38.p14 primary assembly FASTA (3.1 GB), and the Ensembl 114 / GENCODE-v46 GTF (1.7 GB). VX was built with LDC 1.41 in release-aggressive mode (`-O3`, `-mcpu=znver2`, `-boundscheck=off`, AVX2/FMA/BMI2 feature targeting); all timings are warm-cache medians across three to five replicates.

#### Load time (internal track-registration)

After VX is running, a `load_file` request for a new track completes in 0.016 s [0.014, 0.027] for FASTA, 1.17 s [1.13, 1.21] for GTF, and 0.42 s [0.38, 0.63] for BAM. These numbers reflect VX’s design choice to memory-map FASTA, reuse a pre-built `.gidx` for GTF, and open only the BAM index at load time, deferring read fetches to the first viewport draw.

#### Cold start (process-launch to first populated snapshot)

A fair apples-to-apples comparison with IGV is the wall-clock from launching the application to the first snapshot of a chosen viewport after loading the scenario’s inputs, the latency a user experiences when they open a session. For FASTA-only (L1), VX release takes 3.50 s [3.50, 3.68] vs IGV 11.42 s [11.42, 11.72], a 3.3-fold reduction driven by VX’s native D startup and memory-mapped FASTA rather than IGV’s JVM-plus-`.fai`-load path.

#### Memory (steady-state RSS at the same viewport)

Resident set size for VX grew monotonically from 7.7 GB with the FASTA alone (L1) to 10.3 GB after adding the GTF (L2), 12.8 GB with one BAM (L3), 13.8 GB with five (L4), 15.2 GB with ten (L5), and stabilised at 15.2 GB with twenty (L6). IGV at the same scenarios used 693, 629, 728, 1203 and 1243 MB respectively (one to two orders of magnitude below VX). The gap is a deliberate design trade-off: VX memory-maps the entire reference and eagerly parses the GTF into its `.gidx`, paying up-front to deliver sub-second track registration and per-viewport random access; IGV reads both lazily through the `.fai` and its own `.idx`, keeping the footprint small but re-paying the random-access cost on every seek.

#### Rendering

The screenshot endpoint measures GL render → PNG encode → HTTP return for a 1280-pixel viewport; IGV has no comparable scriptable surface, so this metric is internal to VX. VX release sustains 36.1 fps at L2, 35.6 fps at L3, 36.4 fps at L4, 36.6 fps at L5 and 36.6 fps at L6, with round-trip times at 23 ms across the board, indicating the path is bounded by PNG encode plus HTTP transport rather than by per-track rendering cost. Interactive panning and zooming are faster still, because the encode+transport overhead is absent.

#### Analysis runtime

GCContent ran in 0.031 s at a 1 Mb window, 0.523 s at 10 Mb, 0.523 s across chr19, and 0.521 s genome-wide (essentially constant), as the sliding-window loop vectorises cleanly and is bounded by sequence streaming rather than compute. PerExonCoverage took 14.2 s for a 1 Mb window, 13.0 s for 10 Mb, 12.7 s for chr19, and 1 469.5 s genome-wide (two repeats; 907.7 s cold followed by 2 031.4 s after the 23 GB BAM had churned the page cache, confirming the genome-scope wall-time is dominated by BAM random-access I/O rather than compute). SimplePeakCalling scaled from 9.0 s (1 Mb) to 101.5 s (10 Mb) to 307.6 s (chr19); MappingQualityDistribution took 0.53 s (1 Mb), 1.58 s (10 Mb), and 8.14 s (chr19). Both SimplePeakCalling and MappingQualityDistribution at genome scope were omitted from this bench: projected wall time per repeat exceeded the host’s 30 GiB envelope, and the same skip set is retained in the published `bench_analysis_runtime_release.py` driver. Both are recoverable on a 64 GiB host or after the per-contig streaming improvements. Relative to an earlier debug-build pass on the same inputs, the release-aggressive rebuild tightens GCContent by roughly an order of magnitude at all scopes, PerExonCoverage and SimplePeakCalling by ≈ 1.7–2.2*×* at region and chromosome scopes, and leaves the genome-scope PerExonCoverage cost I/O-bound as noted above.

#### Caveats

Three points of context apply to these numbers. First, they are single-host, warm-cache medians: BAM indices were cached after first touch and no kernel page-cache drop was performed between runs. Second, VX render throughput is measured through the screenshot endpoint and is therefore an upper bound on, rather than a direct measure of, interactive responsiveness; IGV exposes no matching scriptable surface for a like-for-like comparison. Third, we restrict the head-to-head to VX versus IGV (Figure 3): adding JBrowse 2 would introduce a web-hosted comparator whose server-round-trip cost is not directly comparable to local desktop wall-clocks.

**Figure 3:**
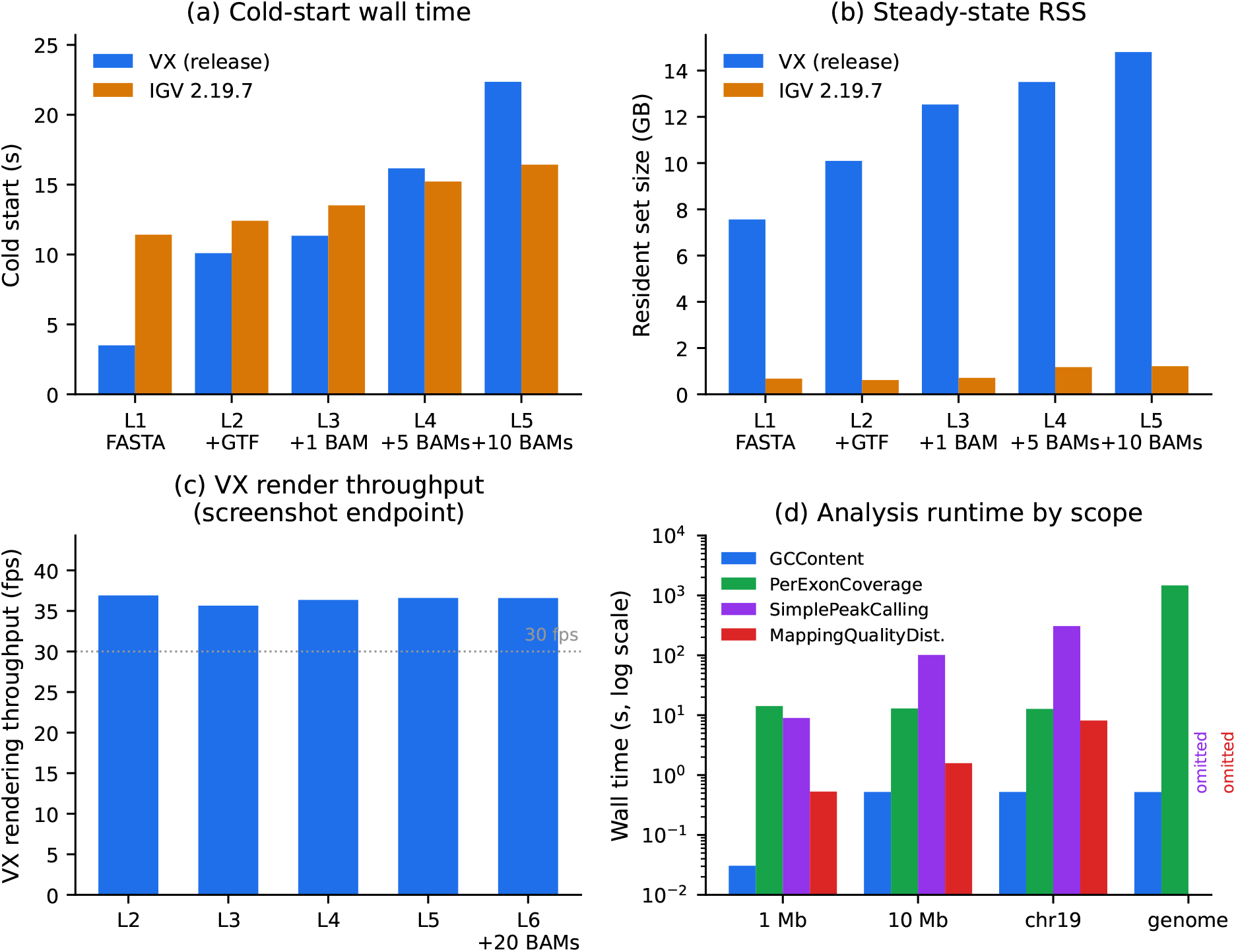
Performance benchmarks on an AMD Ryzen 5 4600G / 30 GiB workstation, VX built LDC 1.41 release-aggressive (`-O3 -mcpu=znver2 -boundscheck=off`) versus IGV 2.19.7 (Java 21). Inputs: full ENCFF754JEN BAM (3.15 *×* 10^8^ reads, 23.2 GB), GRCh38.p14 FASTA (3.1 GB), Ensembl 114 GTF (1.7 GB). **(a)** Cold start (process-launch to first populated snapshot), scenarios L1 (FASTA), L2 (+GTF), L3–L5 (+1, 5, 10 BAMs). **(b)** Steady-state resident set size at the same scenarios. **(c)** VX rendering throughput via the screenshot endpoint (IGV has no comparable scriptable surface). **(d)** VX analysis runtime across scope (1 Mb / 10 Mb / chr19 / genome); `genome-scope SimplePeakCalling and MappingQualityDistribution` omitted as their projected wall time exceeded the host memory envelope. All values are warm-cache medians over 3–5 replicates.

### 3.4 Case study 1: Agent-driven short-read RNA-seq

To demonstrate end-to-end agent control at genome scale, we recorded a live session in which Claude (via the MCP server) was asked to explore an unannotated alignment, identify highly expressed regions, and quantify per-gene expression. The inputs were ENCODE ENCSR000AEM (K562 polyA+ RNA-seq, replicate ENCFF754JEN) subsetted to chromosome 19 (11.07 M reads, 667 MB), paired with the matching chr19 GENCODE annotation and FASTA.

From four user turns the agent issued more than thirty HTTP/MCP calls. Discovery ran entirely inside VX: `SimplePeakCalling` on the alignment track produced a BED of 81 peaks, auto-attached as a Feature track, and four regions were captured at 2*×* scale via the PBO screenshot endpoint. Quantification used `PerExonCoverage` over the alignment and Gene tracks, writing a 14-column TSV covering all 97 829 chr19 exons in 22.2 s. Gene-level aggregation placed FTL, RPS11, RPL13A, SNORD33 and RPS5 at the top. These are canonical ribosomal-protein and translation-associated loci which are expected to dominate K562 polyA+ expression, hence a biologically coherent result. Figure 4 shows the endstate viewport at chr19:49.484–49.502 Mb: the FLT3LG / RPL13A / RPS11 cluster, a full splice-junction ladder, and the Peaks track stacked below.

**Figure 4:**
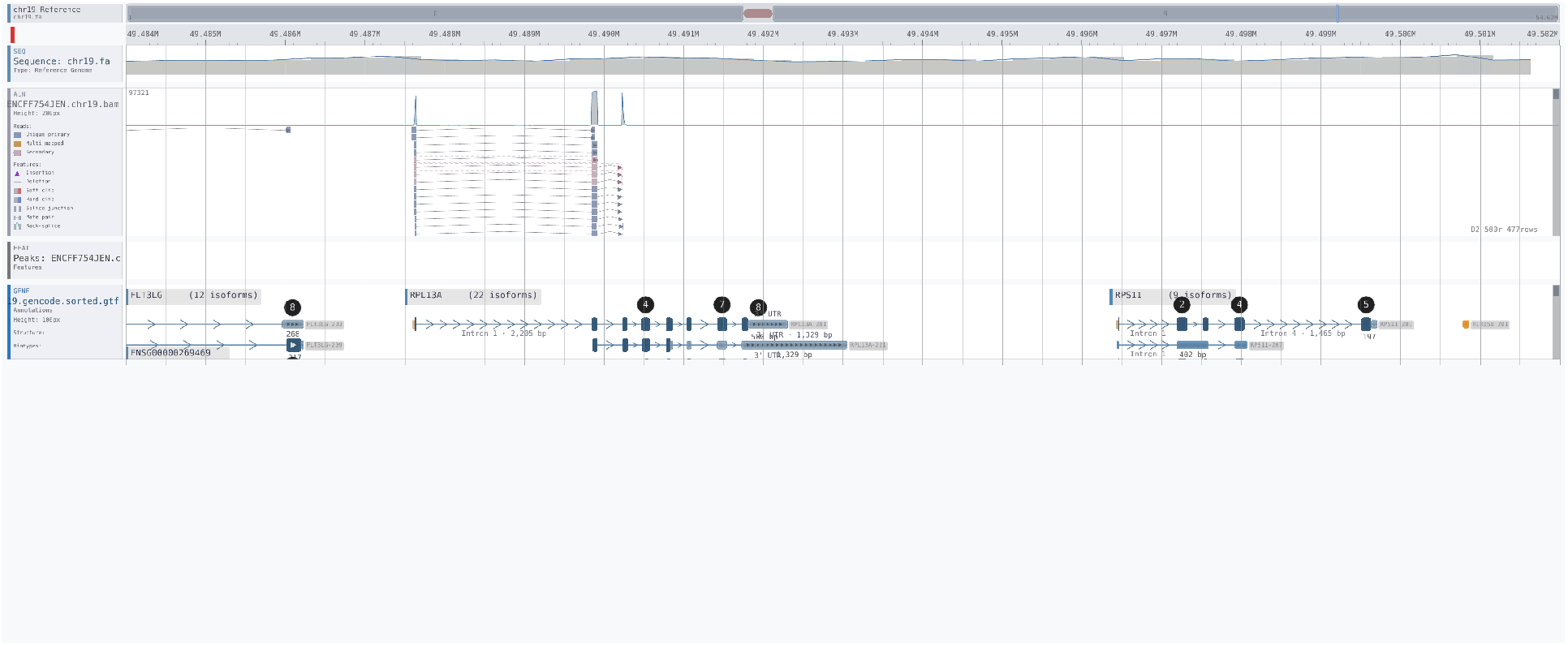
Case study 1 end-state: agent-driven short-read RNA-seq on human GRCh38 (ENCODE ENCSR000AEM, K562 polyA+, primary replicate ENCFF754JEN, chr19 subset). The agent (Claude) loaded the chr19 reference, GENCODE chr19 annotation, and the chr19-subset alignments; ran `SimplePeakCalling` (81 peaks) and `PerExonCoverage` (97 829 exons, 22 s wall clock) via the MCP-wrapped analysis framework; and navigated to the most highly-expressed locus on chr19, the *FLT3LG*/*RPL13A*/*RPS11* ribosomal-protein cluster at chr19:49.484–49.502 Mb. Tracks (top to bottom): Sequence, Alignment (with live coverage histogram, splice junctions as arcs, multi-mapped reads in orange), Peaks (auto-attached Feature track from `SimplePeakCalling`), and Gene annotation (22, 22, and 6 annotated isoforms respectively). Full session transcript in S3.

The session also surfaced internal inconsistencies: the peak caller and `BinStatistics` used a different BAM pre-load path from the quantification analyses, silently truncating genome-scope runs, and several responses lacked a `scope_applied` acknowledgement. These were fixed before the §3.3 benchmarks; the expression table was unaffected. The case illustrates the intended workflow: the researcher specifies intent, the agent drives VX through the same interface a human would use, and the session yields figures and tidy result files for downstream analysis.

### 3.5 Case study 2: Variant inspection in a high-confidence benchmark region

Case 2 exercises the variant-inspection workflow against a community gold-standard truth set. The inputs are the GIAB HG002 NISTv4.2.1 small-variant benchmark restricted to GRCh38 chr21: the high-confidence VCF (56 116 variants), the matching `_noinconsistent.bed` confidence intervals, the chr-prefixed Ensembl release 114 annotation, the chr21 reference FASTA, and a 2.5 GB chr21 slice (11.0 M mapped reads) of the GIAB 2 *×* 250 bp Illumina alignment, sliced from the full 130.8 GB WGS BAM via remote `samtools view -b URL chr21` against the GIAB FTP. All five files were loaded through `vx_load_file`, producing a single chr21 reference group with sequence, gene, alignment, variant and confidence-region tracks stacked in load order.

To exercise reproducible figure production we then drove the viewport loupe entirely from the API. A new `set_magnifier(active, genome_x, …)` command (issue B20, ∼ 80 LOC across `main_window.d` and the MCP wrapper) maps a genomic position to widget pixels via the current viewport, overrides the `Alt`+hover gating for unattended capture, and queues a redraw; the same field set used by the keyboard path is reused so manual and scripted magnifier states are indistinguishable. Figure 5 was then produced unattended: navigate to chr21:35,885,200–35,885,700, `set_magnifier` centred on chr21:35,885,455 at 3 *×* zoom, screenshot via the PBO viewport endpoint at 2 *×* scale. The captured frame shows a heterozygous 21 bp deletion (ATAATGATAAACTGGATAAAG → `A`, allele depth 271/210) as a clean stairstep in the read pile-up, the corresponding benchmark VCF marker pinned beneath, the surrounding `_noinconsistent` confidence interval as a continuous Feature bar, and the magnified detail of the breakpoint inside the loupe popup.

**Figure 5:**
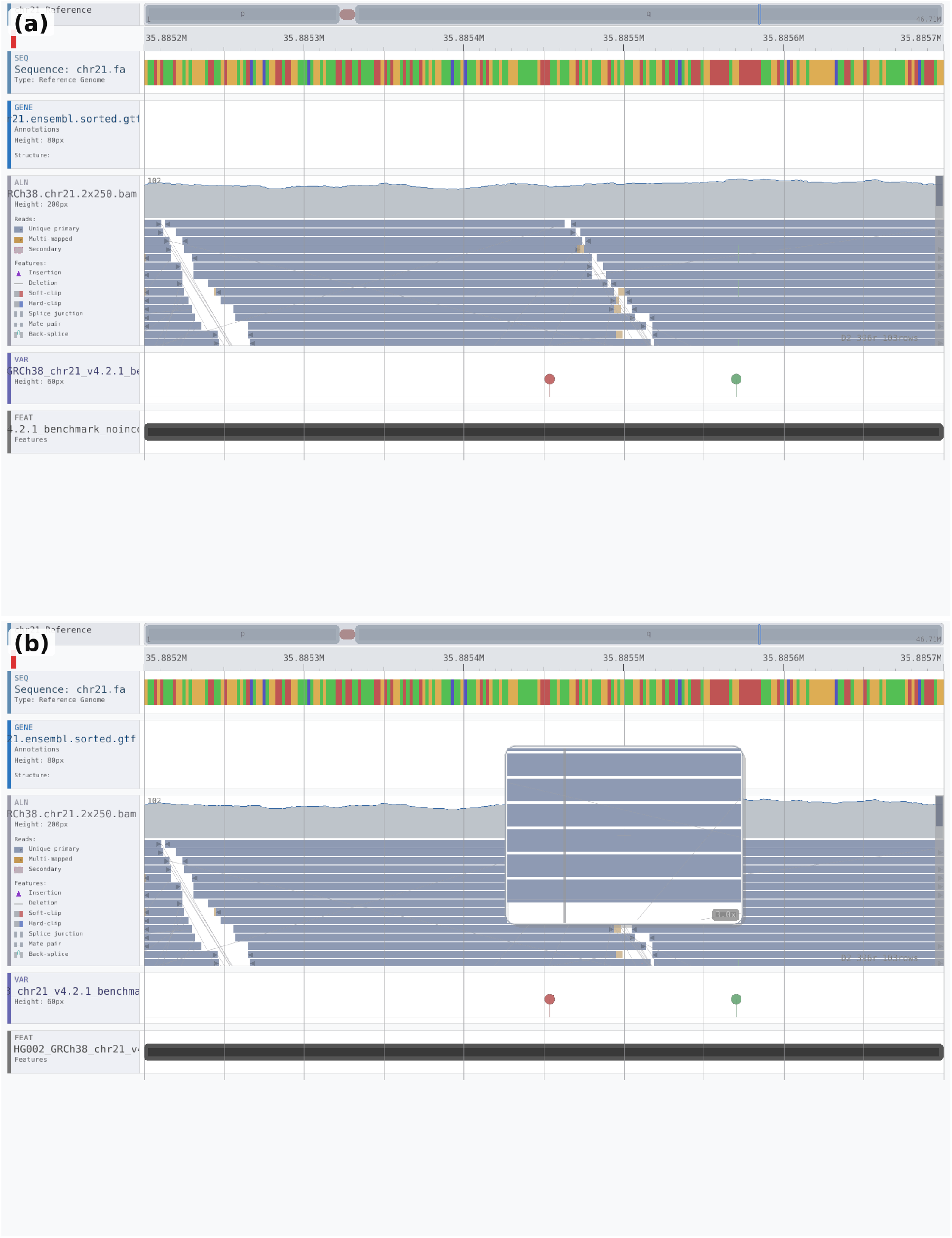
Case study 2 end-state: interactive variant review on GIAB HG002 chr21 (NISTv4.2.1 high-confidence small-variant benchmark; chr21 slice of the 2 *×* 250 bp Illumina alignment, 11.0 M reads). Both panels are captured at chr21:35,885,200–35,885,700; tracks from top: Sequence, Gene (Ensembl r114), Alignment+coverage, Variant (GIAB VCF), Feature (_noinconsistent confidence bar). **(a)** Overview of a heterozygous 21 bp deletion ATAATGATAAACTGGATAAAG → A (AD 271/210), visible as a clean stairstep in the read pile-up, with the benchmark VCF marker pinned beneath. **(b)** The same frame with the API-driven magnifier loupe active (`set_magnifier` centred on chr21:35,885,455, 3 *×* zoom): the breakpoint is resolved to single-base detail without leaving the viewport. Both frames were produced unattended via the MCP server.

### 3.6 Case study 3: Transcriptome browsing and base modifications (two panels)

Case study 3 exercises VX on direct RNA sequencing (DRS) data from the Oxford Nanopore SQK-RNA004 chemistry, where each read carries per-base modification calls (m^6^A at DRACH motifs, 5mC, and pseudouridine) as MM/ML SAM tags [Oxford Nanopore Technologies]. We chose two deposits: (3a) a 10 000-read human pool (HEK293T, HeLa, HepG2, K562; Zenodo 15011476, 365 MB POD5, CC BY) [Mangiante and Ibberson, 2025], and (3b) a 77 527-read *Saccharomyces cerevisiae* aa-tRNA-seq sample (ENA PRJEB90828/ERR15278680, `S288C_deac_SC_rep3`, 1.93 GB POD5 archive) [White and Others, 2025]. Both POD5s were basecalled on the Kaya HPC facility with Dorado 1.3.0 using the RNA004 sup model at v5.1.0 and three modification models in a single pass (m6A_DRACH@v1, pseU@v1, m5C@v1); the human sample additionally used `–estimate-poly-a`, yielding poly-A tail length estimates for 9 703 of 10 088 reads (mean 88 nt). Reads were aligned with `minimap2 2.28`: `-ax map-ont -uf -y -k14` against the Ensembl r114 human cDNA reference for 3a (giving native transcript-coordinate alignments), and `-ax map-ont -uf -y -k10 -w5 –no-long-join` against the S288C R64-4-1 genome for 3b (reducing the k-mer seed size to capture the median 59 nt tRNA read length).

Figure 6a shows VX rendering 3a on `ENST00000309268.11` (EEF1A1, 3512 nt), zoomed to the 1–1700 nt covered range, with a single-command navigation (`vx_navigate`) and `vx_set_alignment_options` to enable the per-base modification summary strip and the coloured probability dots on primary reads. The coverage profile plateaus at the viewport cap of 67 reads, the mismatch strip resolves single-nt mismatches at ∼ 1 px pitch, and the ML-threshold filter (`mod_prob_threshold`=0.5) leaves only high-confidence m^6^A (red) and pseudouridine (gold) calls as coloured glyphs inside each read row. Figure 6b shows VX on the3b yeast data at chrXII:823.2–823.7 kb, a tRNA gene cluster near the 3^*′*^ arm of chromosome XII with 719 primary reads in a ∼ 80 nt window, producing the characteristic sharp coverage spike (4515 reads/nt peak) and the dense stack of short (median 59 nt) tRNA-sized reads. The trans-mapped annotation `chrVI:210345` attached to one of the secondary rows illustrates how VX surfaces cross-chromosome split alignments inline. Both panels were produced unattended via the MCP server.

**Figure 6:**
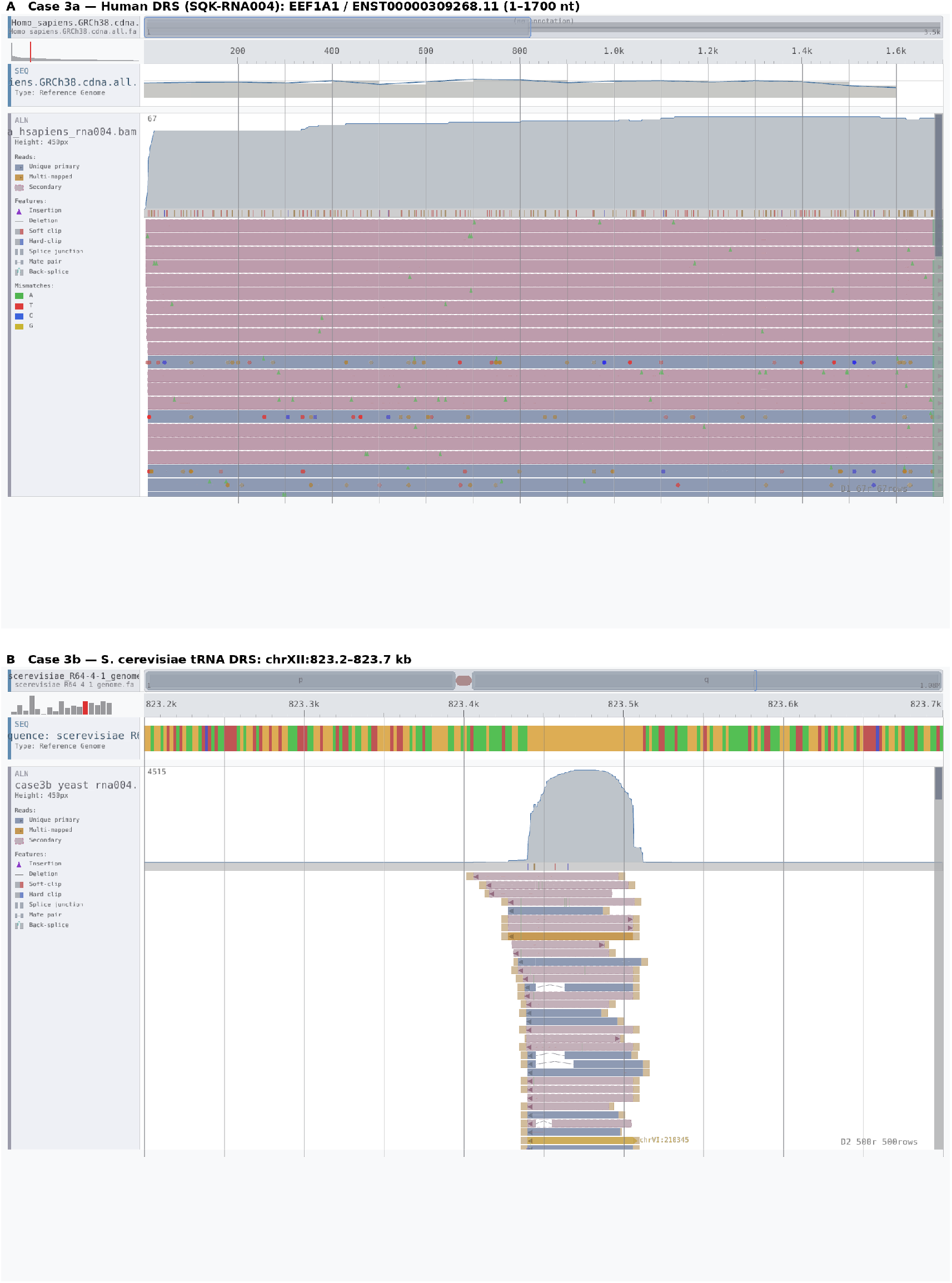
Case study 3 end-state: direct RNA sequencing on two scales. **(a)** Human DRS (SQK-RNA004; Zenodo 15011476, 10 088 reads; basecalled with Dorado 1.3.0 sup v5.1.0 + m^6^A_DRACH/pseU/m5C + `–estimate-poly-a` on Kaya; minimap2 2.28 `-ax map-ont –uf-y -k14` against Ensembl r114 human cDNA). VX view of `ENST00000309268.11` (EEF1A1, 3512 nt; tracks top-to-bottom: Sequence + cDNA annotation bar + zoom strip, Alignment with coverage profile, per-base mismatch summary strip, read stack). `vx_set_alignment_options` has activated the modification summary strip and the coloured probability dots on primary reads (m^6^A red, 5mC blue, pseudouridine gold; ML ≥ 0.5). **(b)** *S. cerevisiae* aa-tRNA-seq (ENA PRJEB90828 ERR15278680, `S288C_deac_SC_rep3`, 81 243 reads; same basecall stack without poly-A; minimap2 `-ax map-ont -uf -y -k10 -w5 –no-long-join` against S288C R64-4-1 genome). VX view of chrXII:823.2–823.7 kb, a tRNA locus with a 4515-read/nt coverage spike over ∼80 nt, showing the median-59 nt tRNA read stack; the `chrVI:210345` annotation on the bottom row is an automatically surfaced trans-mapped secondary alignment. Both panels captured unattended through `vx_navigate, vx_set_alignment_options`, and `vx_screenshot_track` via the MCP server.

### 3.7 Case study 4: Long-read PacBio HiFi in *A. thaliana*

Case study 4 exercises VX on long-read DNA in a non-animal genome. The inputs are ENA PRJEB46164 run ERR6210723 (*A. thaliana* PacBio HiFi), aligned against the Col-CEN v1.2 telomere-to-telomere reference with `minimap2 -ax map-hifi` on the Kaya HPC facility (16 cores, 267 s wall clock), paired with the Araport11 gene-model lift and a self-called VCF from DeepVariant and Sniffles. Files were loaded through `vx_load_file`, producing a single Col-CEN reference group with Sequence, Gene, ATHILA repeat, Alignment, and Variant tracks stacked in load order.

Figure 7 shows the end-state viewport at Chr1:13–18 Mb, which covers the Col-CEN-resolved Chr1 centromere: 129 607 HiFi reads in the window at∼ 4*×* coverage, 25 ATHILA repeat copies (the centromere retrotransposon family re-resolved by the Col-CEN assembly), and a mean read length of ∼13 kb. The viewport was driven through `vx_navigate`, layout through `vx_set_track_height`, and the screenshot captured via `vx_screenshot_track`, all via the MCP server. This case confirms that the same contracts used for short-read RNA-seq (Case 1), variant review (Case 2), and direct RNA sequencing (Case 3) operate without modification on long-read DNA in a plant genome, and that VX renders centromeric repeat stacks and ∼13 kb reads without client-side adaptation.

**Figure 7:**
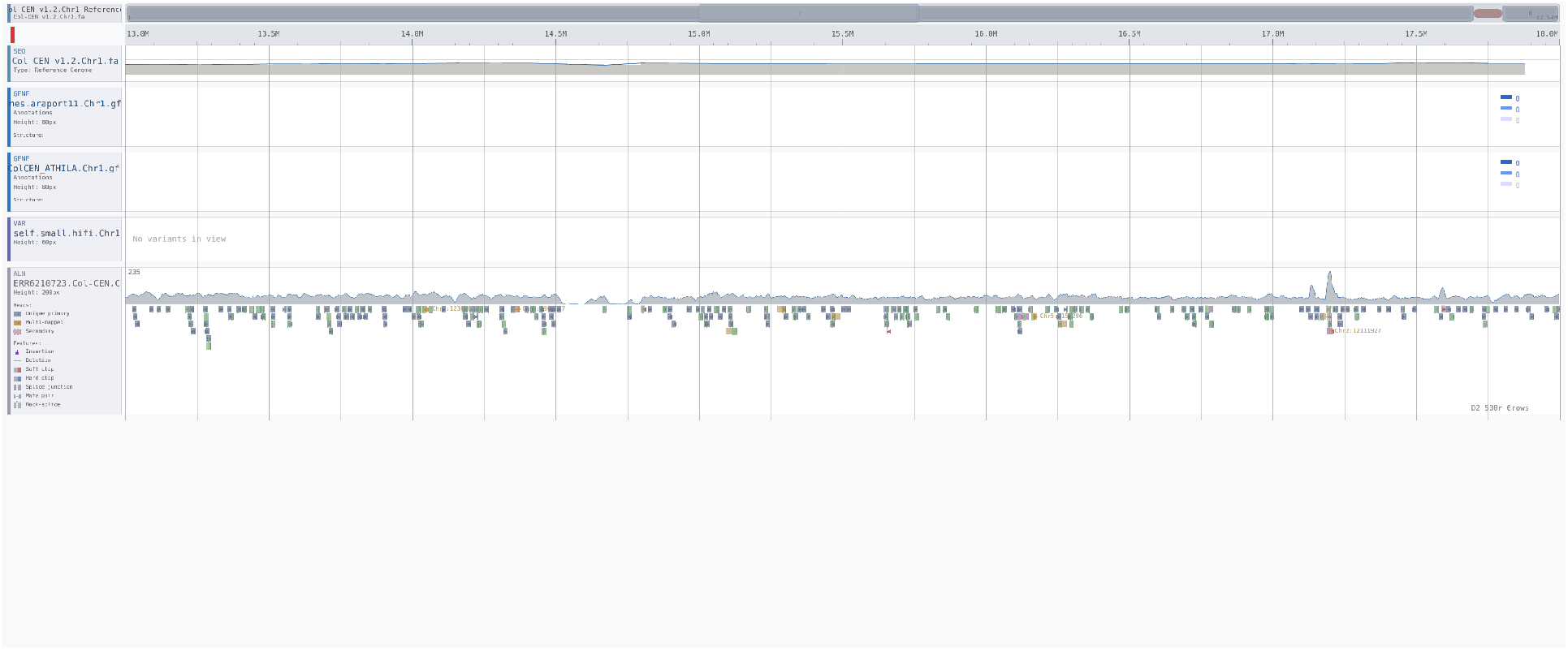
Case study 4 end-state: long-read PacBio HiFi alignment on *Arabidopsis thaliana* Col-CEN v1.2 (PRJEB46164, run ERR6210723). Viewport at Chr1:13–18 Mb covers the Col-CEN-resolved Chr1 centromere (129 607 HiFi reads in this window, ∼ 4*×* coverage from a single sequencing run). Tracks (top to bottom): Sequence (Col-CEN v1.2 reference), Gene (Araport11 lift), ATHILA repeat elements (25 annotated copies in this window, the Col-CEN re-resolved centromere repeat family), Alignment with HiFi-length reads (mean read length ∼ 13 kb), and Variant (DeepVariant + Sniffles self-VCF). Reads were aligned on the Kaya HPC facility with `minimap2 2.28` (`-ax map-hifi`), 16 cores, 267 s wall clock. Capture driven from VX via the MCP server.

## 4 Discussion

### 4.1 Design rationale

VX is deliberately a desktop application rather than a web service. Sequencing data is bulky, often subject to data-governance constraints, and is typically already on the researcher’s machine or an institutional mount; routing it through a browser session adds latency, memory pressure, and transit cost without a clear benefit for the single-user, single-dataset case we target. Direct file access also allows VX to use the native random-access structures of FASTA, BAM, bigWig, and bigBed without an intermediate server-side adaptation layer. Rendering is performed in OpenGL so that the cost of drawing many tracks, or of the magnifier’s sub-region re-projection, is paid on hardware built for exactly that task.

The choice of D follows the same logic. VX requires C-level control of memory layout, straightforward interoperability with C libraries (FreeType, `ffmpeg`), and a garbage collector that keeps the high-level code tractable; D supplies all three in a single toolchain [D Language Foundation]. Finally, the agent-control layer is not a cosmetic addition. Large-language-model assistants are now a routine component of day-to-day bioinformatics work [Thirunavukarasu et al., 2023], and their usefulness is bounded by whether they can observe the same state as the human user and act through the same interface. The HTTP API and MCP server make that boundary conductive rather than opaque, without demanding that the human give up a responsive GUI.

### 4.2 Source-availability rationale

The VX application is distributed as a freely available binary for non-commercial use, with no material-transfer agreement. The core D codebase is not open-source at this release, primarily for commercial-strategy reasons that the authors intend to revisit in subsequent releases. Importantly, closing the core does not close the interface. The full HTTP API and MCP tool surface are specified in this article and are the same contracts the bundled Python MCP server uses.In practical terms, third parties can therefore write clients in any language, build bindings for Jupyter, Nextflow, or continuous-integration systems, extend or replace the supplied Python shim, and, if they wish, implement a compatible back-end against the same interface specification.

This middle path preserves long-term sustainability for the project while meeting what we consider the substantive requirement of a Software article: that the scientific community can build on, verify, and extend the published work. Reproducibility of analyses performed with VX is supported by the deterministic INI configuration and the documented analysis export formats (BED, BedGraph, TSV), both of which can be archived alongside a study’s primary results independent of the closed core.

### 4.3 Limitations

VX has several known limitations, some by design. Release binaries are currently provided for Linux and Windows; an arm64 macOS `.app` bundle (LDC 1.41.0, `dylibbundler`-bundled dependencies, ad-hoc signed) builds cleanly from the same source tree and passes HTTP-API-level tests, but full GUI validation on physical Apple Silicon hardware. VX operates on a single machine by design and does not include a cloud-native mode: files must be reachable through the local file system or a mounted remote volume, and there is no server-tier component that can be shared between users. The MCP server is a separate Python process, which introduces a runtime dependency beyond the self-contained binary; clients that use only the HTTP API face no such requirement. The embedded HTTP server listens on the loopback interface alone and is intended for local automation, not multi-user access. Finally, VX is an application for exploring and analysing aligned or otherwise processed data; upstream steps (base calling, alignment, variant calling) are out of scope and are expected to be performed by established tools. On the performance side, the §3.3 benchmarks make the current trade-offs explicit: VX is competitive or faster than IGV on interactive rendering and region- and chromosome-scope analyses (GCContent and MappingQualityDistribution complete in under two seconds at 10 Mb, PerExonCoverage and SimplePeakCalling in roughly 10–100 s), but genome-scope analyses over a 23 GB BAM remain I/O-bound and exceed the 30 GiB memory envelope of the test host for `SimplePeakCalling` and `MappingQualityDistribution`. This weakness is recoverable on a larger-memory machine or after the per-contig streaming improvements targeted for the next release. A specific operational limitation is also worth recording: the very first read-fetch that follows a file-open into a previously unvisited region of a large BAM (e.g. our 23 GB ENCFF754JEN test case) currently takes on the order of two minutes on commodity NVMe storage, even though subsequent navigations into the same region complete in well under a second. The cost is dominated by cold BAM-index traversal and first-time pagecache population rather than VX-internal work; the load-status strip (§2.7) makes this cost visible to the user, but reducing it (warm BAM index cache, prefetch of the neighbouring tiles around the first navigation target) is deferred to a future release. Several alignment-rendering refinements (preservation of base-modification tags under subsampling, polyA-tail overlay state, soft-clip indicator state coupling) are scheduled for v0.9.x point releases following the initial public release.

### 4.4 Future work

Four directions are planned or under way. First, a macOS binary to complement the existing Linux and Windows releases, which the current D, GTK 3, and OpenGL stack already supports at the source level. Second, further format coverage: native BCF parsing, CRAM [Hsi-Yang Fritz et al., 2011] support, and long-read-specific rendering refinements for modified-base and poly(A) signal tags produced by current ONT pipelines [Danecek et al., 2021]. Third, expansion of the analysis framework: additional peak-calling variants, differential-expression analyses operating across loaded signal and count tracks, and richer cross-track statistics that build on the existing scope hierarchy. Fourth, a more developed plugin interface, so that domain-specific analyses and track types can be contributed without a core rebuild. Alongside these, three performance-oriented improvements are already scoped from the §3.3 results: per-contig streaming of alignment-derived analyses, which would bring `SimplePeakCalling` and `MappingQualityDistribution` within the 30 GiB envelope at genome scope; a warm BAM read cache shared across analyses within a session, targeting the cold/warm spread observed in genome-scope `PerExonCoverage` (≈ 900 s cold versus ≈ 2 000 s after cache churn); and release-build vectorisation audits of the remaining sliding-window kernels, following the ≈10*×* tightening already observed for `GCContent` under `-O3 -boundscheck=off`.

Two longer-horizon directions are also worth noting. An optional cloud-data module (S3, Azure, and public-bucket access through signed URLs) would let VX open remote objects without manual staging while keeping the desktop-first model intact. A multiuser mode built on shared HTTP-API sessions (effectively a collaborative viewport) would extend the existing agent-control layer and repurpose the same scripting surface for human collaboration.

## 5 Conclusions

VX is a desktop genome and transcriptome viewer designed for the AI-assisted era. By exposing its full functionality through an embedded HTTP API and a Model Context Protocol server, it allows agents and scripts to drive exploration, run analyses, and produce figures through the same interface that serves the human user, while a responsive OpenGL-rendered GUI, a magnifier popup, and an integrated analysis framework with an explicit scope hierarchy support direct interactive work. VX handles chromosome- and transcript-coordinate data uniformly, filling a gap that existing browsers address only through custom references or plugins. We see VX as complementary to IGV, JBrowse 2, the UCSC Browser, and pyGenomeTracks rather than a replacement: it serves the niche these tools were not designed for: programmable, agent-accessible desktop visualisation with an integrated analysis layer. The binary is freely available for non-commercial use; the HTTP API and MCP protocol are fully specified in this article, so the scientific community can build on, verify, and extend the published work independently of the core D implementation.

## Availability and requirements

- **Project name:** VX Genome Viewer
- **Project home page:** https://github.com/Arnaroo/VX
- **Archived version:** Permanently archived on Zenodo with concept DOI 10.5281/zenodo.20234450, which always resolves to the latest archived release; the snapshot accompanying this submission is version DOI 10.5281/zenodo.20234451 (release tag `v0.9.0.1`, archival re-publish of the `v0.9.0` “Aardvark” public pre-release). VX Linux and Windows release binaries in the `bin/Dist` distribution (`vx-linux`, `vx-windows`); Python MCP server source accompanying this article.
- **Operating system(s):** Linux (tested on Manjaro with a modern kernel) and Windows release binaries are provided; an arm64 macOS `.app` bundle and `.dmg` also build cleanly from the same source tree and pass HTTP-API smoke tests, with full GUI verification on physical Apple Silicon hardware pending (see §4.3)
- **Programming language:** D (dlang), with the Model Context Protocol server implemented in Python 3
- **Other requirements:**
  – GTK 3 (≥ 3.22) with GtkGLArea support
  – An OpenGL 3.3-capable driver
  – FreeType 2
  – `ffmpeg` ≥ 4.0 on `$PATH` (required only for viewport video recording)
  – Python ≥ 3.10 with `httpx` and `fastmcp` (required only for the bundled MCP server)
- **Licence:** Proprietary; freely available for non-commercial use. The HTTP API specification and the MCP tool reference are published under a permissive documentation licence to permit independent client implementations.
- **Any restrictions to use by non-academics:** Commercial use requires a separate licence, obtainable from the corresponding authors.

## Acknowledgements

The authors are very grateful to the Biocodecs Group, the RNA Innovation Foundry (RIF), and The Australian Centre for RNA Therapeutics in Cancer (ACRTC) teams for useful and enabling discussions throughout the project. The authors would like to acknowledge their use of the Kaya HPC facility at UWA and the unwavering, highly enabling support provided by the Kaya HPC team, including the amazing contributions and work of Chris Bording. The authors also thank an external pre-release user whose stress-testing on a 23 GB short-read alignment dataset directly motivated the responsive-feedback design described in §2.7. Use of large-language-model assistants during development and manuscript preparation is described in §2.9.

## Authors’ contributions

N.E.S. led conceptualisation, initial software design and implementation, and drafted the original manuscript. N.E.S. and A.C. jointly carried out methodology, validation, formal analysis, investigation, data curation, visualisation, manuscript review and editing, supervision, project administration, and funding acquisition. Both authors read and approved the final manuscript.

